# The predictive potential of key adaptation parameters and proxy fitness traits between benign and stressful thermal environments

**DOI:** 10.1101/2021.04.29.441345

**Authors:** Jennifer M. Cocciardi, Eleanor K. O’Brien, Conrad J. Hoskin, Henry Stoetzel, Megan Higgie

## Abstract

Understanding the adaptive potential of a species is key when predicting whether a species can contend with climate change. Adaptive capacity depends on the amount of genetic variation within a population for relevant traits. However, genetic variation changes in different environments, making it difficult to predict whether a trait will respond to selection when not measured directly in that environment. Here, we investigated how genetic variances, and phenotypic and genetic covariances, between a fitness trait and morphological traits changed between thermal environments in two closely-related *Drosophila*. If morphological traits strongly correlate with fitness, they may provide an easy-to-measure proxy of fitness to aid in understanding adaptation potential. We used a parent-offspring quantitative genetic design to test the effect of a benign (23°C) and stressful (28°C) thermal environment on genetic variances of fecundity and wing size and shape, and their phenotypic and genetic covariances. We found genetic variances were higher within the stressful environment for fecundity but lower within the stressful environment for wing size. We did not find evidence for significant phenotypic correlations. Phenotypic and genetic correlations did not reveal a consistent pattern between thermal environments *or* within or between species. This corroborates previous research and reiterates that conclusions drawn in one environment about the adaptive potential of a trait, and the relationship of that trait with fitness, cannot be extrapolated to other environments *or* within or between closely-related species. This confirms that researchers should use caution when generalising findings across environments in terms of genetic variation and adaptive potential.

## Introduction

Climate change is causing increased temperatures that will impose stress on species (Thomas et al. 2004). Many species lack the ability to disperse to more optimal environments (Bellard et al. 2012; Ceballos et al. 2017), and will have to adapt to the stressful temperatures to survive in the long-term (Thomas et al. 2004; Hoffmann and Sgrò 2011). Adaptation potential will depend on the amount of genetic variation in traits relevant to the selection imposed by environmental change (Fisher 1930; Falconer and Mackay 1996), and adaptation will need to be rapid given the speed of human-induced climate change. Understanding the adaptive potential of species, especially those currently living close to their upper thermal limits, is therefore crucial in today’s changing climate (Urban et al. 2016; Funk et al. 2019; Shaw 2019).

Importantly, genetic variation is context-dependent — meaning the amount of genetic variation in a trait in a given population can change under different environments (Falconer and Mackay 1996; Hoffmann and Schiffer 1998; Sgrò and Hoffmann 1998a, c; Hoffmann and Merilä 1999). Short-term environmental changes can play an important role in adaptive evolution (Wood and Brodie 2016) and can induce a similar or larger change in genetic variance than changes to the genetic architecture that accumulate over hundreds of generations between populations (for review, see Wood and Brodie 2015). Increases in environmental variability, such as those predicted with climate change, will therefore directly affect the rate of evolution of a trait — as environments get warmer, not only may the type of selective pressure change, but also the potential for the trait to respond to selection. This is important because as researchers aim to determine whether species can adapt to climate change, the changing climate itself may increase or decrease adaptation potential.

Much research has focused on examining whether there is a consistent pattern to changes in expression of genetic variance (for reviews, see Sgrò and Hoffmann 2004; Rowiński and Rogell 2017; Fischer et al. 2020). However, there is no consensus on whether stressful conditions increase or decrease the expression of genetic variance. The majority of studies focus on quantifying genetic variance by calculating heritability (*h^2^*), which describes the relative amount of genetic variance due to additive effects standardised by total phenotypic variance. Heritability can then be used to predict the magnitude of the response to selection via the breeder’s equation. These studies show both increases (e.g., Sgrò and Hoffmann 1998a, c; Swindell and Bouzat 2006) and decreases (e.g., Hoffmann and Schiffer 1998; Bubliy et al. 2001; Kristensen et al. 2015) in heritability under stressful conditions. Increased heritability may result from novel genetic variance that is expressed when exposed to new conditions (i.e., ‘cryptic genetic variance’; see review by Hoffmann and Schiffer 1998; but also see Swindell and Bouzat 2006). Decreased heritability may result from low cross-environment genetic covariances (Fischer et al. 2020), or environmental variance increasing while other variance components remain the same (for example see Hoffmann and Schiffer 1998). Recently, studies have recommended quantifying genetic variance using parameters standardized by the trait mean — such as coefficient of additive variance (*CV_A_*) and its square, evolvability (*I_A_*) — because estimates of heritability, which are standardised by the phenotypic variance, can be influenced by sources of non-genetic environmental variation that may preclude comparison across environments and traits (Houle 1992).

Assessing the effect of a changing environment on genetic variance is further complicated when attempting to measure genetic variance across different environments for fitness. Direct fitness (reproductive success) is often difficult to measure in the wild because of uncontrolled and unmeasured factors (Orr 2009), and in the laboratory due to time and logistical constraints (Rosenberg 1982; Nguyen and Moehring 2015). Instead, a morphological trait that strongly correlates with fitness, and is more easily measured, may provide a good proxy when fitness measures are difficult to obtain. If phenotypic correlations between a morphological trait and fitness are strong, researchers can use the easier-to-measure trait to predict genetic variation of fitness across different environments (Arnold 1983).

More importantly, a strong phenotypic correlation may indicate that two traits are genetically linked through physical linkage, pleiotropy, or linkage disequilibrium (Cheverud 1988; Conner and Via 1992; Roff 1995; Blows and Hoffmann 2005). This means a positive genetic correlation between traits could aid adaptation to novel environments if selection favours that trait combination through augmenting the effect of selection on the correlated fitness trait (Blows and Hoffmann 2005; Agrawal and Stinchcombe 2009; Walsh and Blows 2009; Holman and Jacomb 2017). Therefore, determining genetic correlations of traits with fitness is an important part of the puzzle when predicting evolutionary potential.

However, much like genetic variation in individual traits, phenotypic and genetic covariances between traits (or between a trait and fitness) can vary depending upon the environment in which they are measured (Sgrò and Hoffmann 2004) — meaning measurements obtained in one environment cannot necessarily be generalized to other environments. For example, in a beetle, adult female body mass and her egg size were positively correlated on one host-plant species and negatively correlated on a different host-plant species (Czesak and Fox 2003). Changes to genetic correlations can result within novel environments due to genotype-environment interactions — where genes that affect a trait in one environment may not be influential in a different environment (Sgrò and Hoffmann 2004). In some instances, the loci that contribute to covariances through pleiotropy or physical linkage have specifically been found to be influenced by environmental effects (e.g., Hausmann et al. 2005; Gutteling et al. 2007). However, more empirical data are needed to understand whether there are patterns to how genetic variances and covariances of morphological and fitness traits vary across thermal environments (Rowiński and Rogell 2017; Fischer et al. 2020).

*Drosophila* are often used to investigate genetic variances due to their short generation time and ability to produce large numbers of offspring that allow for quantitative genetic experimental designs. Fecundity is a commonly assessed fitness trait in *Drosophila*. However, measuring fecundity can often prove time- and labour-intensive and logistically challenging. Ecological theory assumes that body size is correlated with fecundity, with larger individuals exhibiting a higher fecundity (Chiang and Hodson 1950; Santos et al. 1992; Robertson 1956), and wing length has been shown to phenotypically correlate with fecundity (Tantawy and Vetukhiv 1960; Woods et al. 2002). However, two key studies examining the relationship of wing length and fecundity in *Drosophila* when exposed to stressful environments found mixed evidence. Sgró and Hoffmann (1998a) did not detect a significant positive phenotypic or genetic correlation in a cold-stress, heat-stress, or benign environment. They also did not find a significant genetic cross-environment correlation (parents raised in one environment and offspring raised in a different environment) between cold-stress, heat-stress, or benign environments (Sgrò and Hoffmann 1998a) — meaning that they did not find a correlation between wing length and fecundity among and between any experimental environment. Conversely, Woods et al. (2002) found significant positive phenotypic correlations (for two of three generations) and significant positive genetic correlations between wing length and fecundity in a stressful environment, but not in a benign environment.

With advances in technology over the past decade (i.e., advances in microscopic imaging and digitizing), more intricate morphological traits such as wing size and wing shape have been increasingly used in place of wing length. However, very few studies have examined genetic variation and heritability in wing size and shape (Gilchrist and Partridge 1999; Hoffmann and Shirriffs 2002; Moraes et al. 2004); and, to our knowledge, only one has examined the phenotypic and genetic correlations of wing size with fecundity (e.g., Woods et al. 2002). Wing size and wing shape in *Drosophila* have a polygenic basis independent of one another (Carreira et al. 2011), so phenotypic and genetic correlations of each of these traits with fecundity may differ. Wing size exhibits a history of directional selection in *Drosophila*, whereas wing shape has been shown to undergo optimizing selection (Gilchrist and Partridge 2001). Although most of the fundamental research uses wing length as a trait that is highly correlated to thorax size (and therefore body size; Chiang and Hodson 1950; Tantawy and Vetukhiv 1960; Santos et al. 1992; Woods et al. 2002), wing size may be a better indicator of overall body size because it is a product of more complex interactions between the different wing compartments (i.e., anterior and posterior compartments; Guerra et al. 1997; Gilchrist and Partridge 1999). Hence, wing size may account for a greater proportion of variation than wing length alone. Wing shape is important for flight performance in *Drosophila* and has been shown to exhibit high heritability (Hoffmann and Shirriffs 2002; Moraes et al. 2004). However, whether selection occurs on wing shape itself, or whether wing shape is correlated with another trait under selection is unknown (Gilchrist and Partridge 2001).

Temperature as a stressor is contextually important in today’s climate, but it has only been used in one *Drosophila* study to assess whether genetic correlations exist between fecundity and wing length, and whether heritability changes between different thermal regimes (i.e., *D. melanogaster*; Sgrò and Hoffmann 1998a with the same data used in Woods et al. 2002). Here, we focused on whether genetic variances in fecundity change across thermal environments, and whether a morphological trait that may be a good proxy of fitness in one environment was also a good proxy in a stressful thermal-environment. We examined the consistency of heritability, coefficient of additive genetic variance, and evolvability between thermal environments (one benign and one stressful), generations, and within and between two sibling species of *Drosophila*. A strength of this study is that we assessed both life history and morphology traits in two closely-related species to see whether this pattern was conserved. We also assessed the phenotypic and genetic covariances of these traits. The correlation of body morphology with fitness informs us about the strength and direction of selection. This is important because patterns of selection in one environment may not reflect similar responses in another environment.

## Methods

### Experimental populations

Two sibling species of fruit fly found along the east coast of Australia were used in this study: *Drosophila serrata*, a generalist species found in forested areas; and *D. birchii*, a specialist species confined to tropical rainforest ecosystems (Schiffer and Mcevey 2006; Higgie and Blows 2008). Mass bred populations from two different geographical areas for each species were used. Each mass bred population was originally created by breeding the offspring of ten isofemale lines collected from field sites within Queensland, Australia. *Drosophila birchii* flies were collected from Paluma National Park (19° 0’16.27”S, 146°12’35.59”E) and Mt. Lewis National Park (16°35’30.36”S, 145°16’27.78”E). *Drosophila serrata* flies were collected from Paluma National Park (19° 0’16.27”S, 146°12’35.59”E) and Raglan Creek (23°42’49.74”S, 150°49’0.10”E). All flies were collected between February and May 2016. Isofemale lines were maintained in controlled laboratory conditions for 18 generations before mass bred populations were created. All stocks were maintained at large population sizes (*N* > 1000) to retain natural genetic variation. Flies were reared on standard *Drosophila* food that contained sugar, yeast, and agar as described in Higgie and Blows (2008). All flies were reared under constantly controlled laboratory conditions of 23°C ± 1°C, 50% relative humidity (RH), and 12 h light:dark cycles.

### Quantitative genetic experimental design

A parent-offspring breeding design was used to assess heritability and phenotypic and genetic covariances of fecundity and wing morphology at a benign (23°C) and a stressful (28°C) temperature (Fig. 1). The benign temperature (23°C) represents an approximate average temperature each species experiences across their range both temporally and spatially, as well as the optimal rearing temperature in the laboratory. A temperature of 28°C was chosen as a stressful thermal environment as it was found to be within the upper margin of the thermal niche for *D. birchii* and to place stress upon *D. serrata* (from pilot studies showing reduced survival). As such, full development was expected in both species at this temperature.

**Figure 1:**
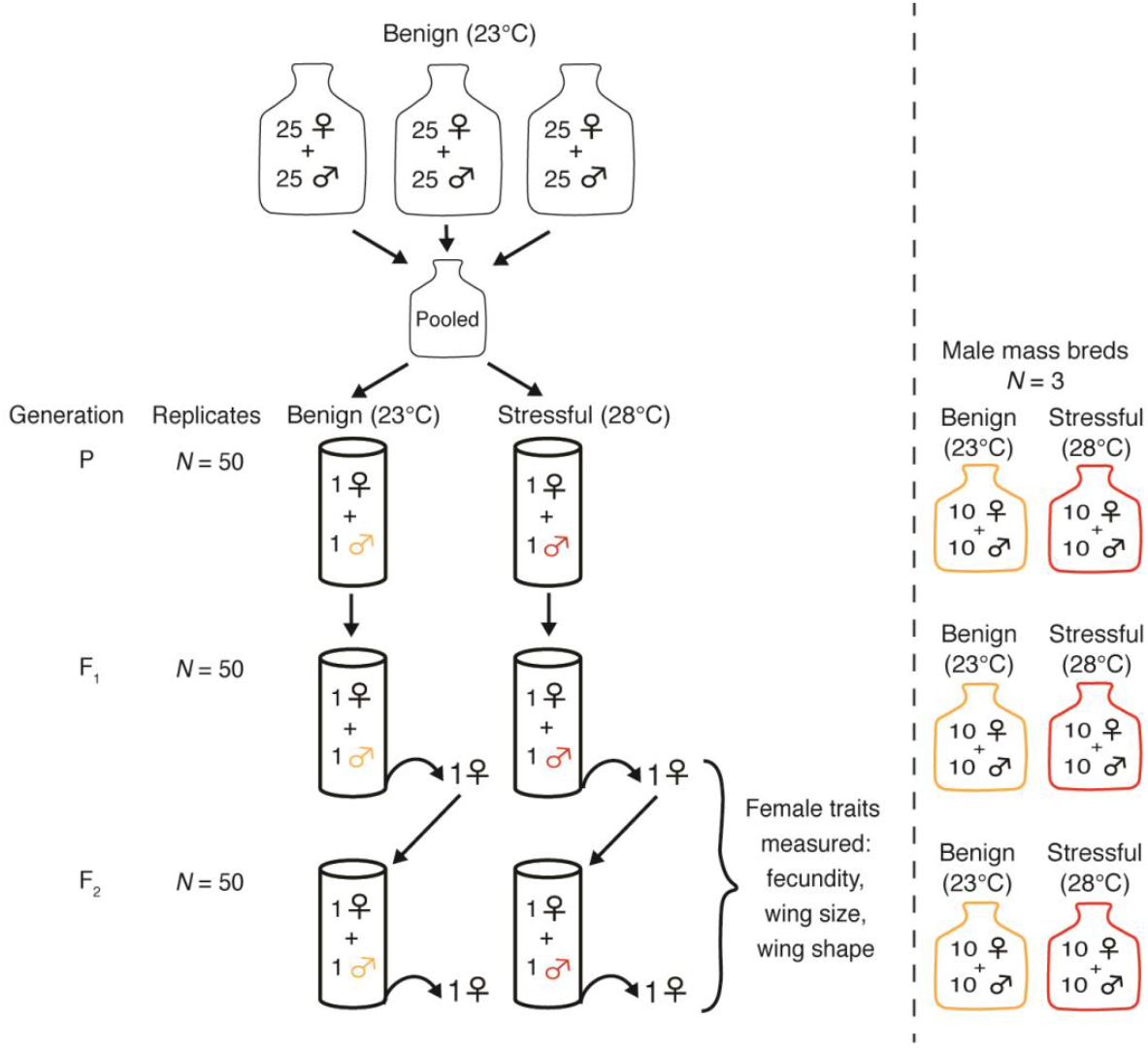
Parent-offspring quantitative genetic experimental design. The parent-offspring quantitative genetic experimental design used to measure female fecundity, wing size, and wing shape on both dams and daughters. This design was used for two populations of both *D. birchii* and *D. serrata*. Experimental female flies were raised in either a non-stressful rearing temperature (23°C) or a stressful temperature (28°C). Mass bred populations were raised alongside each generation and supplied males for mating purposes.

Two generations before the start of the experiment, density-controlled mass bred populations were created for each species and population by sexing 25 virgin females and 25 virgin males from the laboratory stock and placing them in one 300 mL bottle with 100 mL of food. This was repeated three times for each species and population. Flies were removed from each replicate bottle after 72 h and bottles were carded for pupation. Offspring were collected at random and sexed to subsequently create family lines and stock mass bred populations for each species and treatment.

One generation before the start of the experiment (i.e., P generation; Fig. 1), virgin offspring were sexed from the density-controlled mass bred populations using CO2 anesthetization. Flies were placed in 100 mL holding vials with 5 mL of food for 72 h to allow for sexual maturation and full recovery from anesthetization, with 5 individuals per holding vial. After this, one male and one female were randomly collected and placed in a 100 mL glass vial with 10 mL of food, stoppered with a porous stopper, and directly placed in an incubator set to the relevant temperature for each thermal environment treatment. Humidity inside the vials with the stoppers on remained at approximately 90% RH, and a 12 h light:dark cycle was maintained. This was carried out for 50 family replicates for each species, population, and treatment. Mating pairs were allowed to mate for 48 hours before being removed from the vial. This ensured all experimental flies were reared in a controlled and low-density environment. In addition, three low-density stock bottles containing 10 females and 10 males were created and maintained for both the parent and offspring generations to provide a supply of males for mating to assess fecundity (i.e., male mass breds; Fig. 1). Three stock mass bred bottles were maintained in each thermal environment and males were randomly collected from each bottle and mated with a female from the same experimental rearing temperature.

### Fecundity measurements

Virgin female offspring of each family replicate vial were sexed under light anesthetization and placed in holding vials for 72 h. One female (i.e., dam) from each F_1_ family was randomly selected and placed in an empty vial with one virgin male collected from the male stock bottles. Each vial contained a small spoon with 2 mL of food to provide a medium for oviposition. The food was dyed green to aid in counting eggs, and a drop of a live yeast-water solution (1 g baker’s yeast: 10 mL water) was spread over it to promote ovipositing. Vials were immediately placed within their temperature treatment and flies were allowed to mate for 24 h. After 24 h, the spoon was removed and immediately frozen at −19°C for eggs to be counted at a later time, and replaced with a new spoon. This was repeated every 24 h for three days and a cumulative fecundity count was obtained. Cumulative fecundity measurements from the first three days of maturity are significantly correlated with lifetime reproductive success of female *Drosophila* (Pekkala et al. 2011; Nguyen and Moehring 2015). After 72 h, the mating pair was transferred to a rearing vial with 10 mL of food and allowed to mate for the next 48 h period before being removed. Females were then immediately frozen for wing morphology measurements. Daughters of these pairs were collected from each vial and one virgin female offspring from each mating pair was assessed for fecundity and wing traits using the same methods described above.

Fecundity was scored using a microscope and click counter by counting the number of eggs on each spoon. Approximately one quarter of spoons were counted twice, at random, to assess repeatability; a positive correlation close to 1 indicated that counting was highly repeatable between measurements (*r* = 0.994; *P* < 0.001; *N* = 81).

### Wing morphometrics

All dams and daughters were frozen at nine days old to assess wing size and shape. Left wings were removed using fine forceps and mounted on microscope slides with double-sided tape. Wings were photographed using a Leica Image microscope and Leica Application Suite software (LAS v. 3.8). Images were randomized and collated as a TPS file using *tpsUtil* (Rohlf 2010a). Landmarks were placed on ten consistent morphometric wing features of each image (Supplementary Fig. 1) using the program *tpsDig2* (Rohlf 2016). Outliers and landmarking errors were identified using *tpsRelW* (Rohlf 2010b) and corrected or removed before wing measurements were computed.

Landmarked coordinates underwent a Generalized Procrustes Analysis (GPA) superimposition (Rohlf and Slice 1990), where wing size and alignment are adjusted for by superimposing images upon one another over an average configuration. The GPA superimposition has been found to produce estimates with the least amount of error in a study on geomorphometrics (Rohlf 2003). The square root of the summed squared distance between centroid configuration and landmarks is known as the centroid size and provides a measure of overall size (Rohlf and Slice 1990; Rohlf 2000). Although size effects should be removed via the GPA, a correlation between shape and size might still occur (known as allometry), and hence this was also assessed.

In addition to centroid size, the GPA computes a set of Procrustes residuals for each landmark. A principle component analysis (PCA) was conducted upon these to identify variation components, which can be used to describe a single axis of variation in wing shape among individuals (Adams et al. 2004; Zelditch et al. 2004; Gómez et al. 2009). In this instance, the PCA is equivalent to a relative warp analysis because the variation between landmarks was not weighted by bending energy (Zelditch et al. 2004), so PCA scores are equivalent to relative warp (RW) scores. As per common practice, RW scores that explained greater than 5% of variation were used as shape variables (Zelditch et al. 2004; Gómez et al. 2009) to analyse differences in wing shape using the *geomorph* package (Adams et al. 2020) in R.

### Analysis

Data was checked for outliers, homogeneity, normality, and independence as outlined in the protocol described in Zuur et al. (2010) and analyses were performed using the statistical program R (R Core Team 2019). Mean trait values and phenotypic variances (calculated as squared standard errors of the mean trait values) were calculated for fecundity, wing size, and wing shape for both the dam and daughter generations. To test whether thermal environment had an effect on mean trait values and on phenotypic variances, a two-way ANOVA (for thermal environment and generation and its interaction) or a generalized least square model was conducted for each trait and metric, depending on data structure. Population and its interaction with thermal environment was also included when significant. For the multivariate measure of shape (RW score matrix), a permutational MANOVA (also known as a Procrustes ANOVA; Goodall 1991) was used to test for differences in wing shape between thermal treatments, populations, and generations. All analyses were conducted separately for each species.

Narrow sense heritability (*h*^2^) for each trait was calculated from a regression of offspring trait values on maternal trait values (Falconer and Mackay 1996). As we conducted only a single-parent regression, the phenotypic resemblance is equal to half of the genetic variation and thus the slope parameter estimate *β* represents:

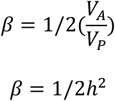

and so, heritability is equal to twice the slope of the regression line (Falconer and Mackay 1996). In parent-offspring regression, estimates can be greatly skewed if variances found in the parental generation and offspring generation differ (Falconer and Mackay 1996). To overcome this, we standardized all traits to a mean of zero and standard deviation of one prior to computation of heritability and genetic covariances (Sgrò and Hoffmann 1998b). The significance of deviations of heritability estimates from zero were assessed using an *F*-test and all *P*-values were adjusted for using the False Discovery Rate method (Benjamini and Hochberg 1995). Standard errors of heritability were obtained directly from the regression model.

To obtain an overall estimate of heritability for wing shape, we followed the equations set forth in Monterio et al. (2002) for estimating heritability from a parent-offspring regression on a multivariate trait (i.e., wing shape with all RW scores included). This was done by first obtaining the coefficient of determination (*R*^2^) from a multivariate linear regression of offspring RW scores onto dam RW scores. The square root of the coefficient of determination (*R*) was then used in the following formula:

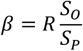

where *β* is the multivariate regression coefficient, *S_O_* is the standard deviation of the offspring trait, and *S_P_* is the standard deviation of the parental trait. The multivariate regression coefficient multiplied by two is then equal to the heritability of the trait (to account for only having half of the genetic variation due to the single-parent-offspring comparison). Significance of deviation from zero for multivariate shape heritability was assessed using a Wilks’ lambda test (Zelditch et al. 2004; Gómez et al. 2009).

In addition, coefficients of genetic variation (*CV_A_*) and evolvabilities (*I_A_*) were calculated following Houle (1992) as:

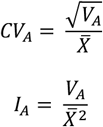

Because we did not directly calculate *V_A_* in this analysis, we obtained estimates based on the method of Garcia-Gonzalez et al. (2012). *V_A_* estimates were calculated by multiplying the total phenotypic variance (*V_P_*) of each trait mean by the narrow-sense heritability (*h*^2^), since *V_A_* = *h*^2^ × *V_P_* (Falconer and Mackay 1996). This is an alternative way to calculate *CV_A_* when researchers do not have the sire variance component (*V_sire_*) or another direct measure of *V_A_* (Garcia-Gonzalez et al. 2012). Standardized data cannot be used to calculate *CV_A_* and *I_A_* because a scaling correction to a zero mean produces a meaningless comparison and undefined value when dividing by the trait mean a second time (Garcia-Gonzalez et al. 2012). The above methods were therefore only performed on non-standardized data and *CV_A_* and *I_A_* values were not calculated for RW scores of wing shape as these are standardized.

The phenotypic correlation among each pair of traits was calculated as the Pearson correlation coefficient. Genetic covariances (*cov_XY_*) were obtained by regressing one trait in the parental generation onto the other trait in the offspring generation, in both directions (Supplementary Table 5), adjusting for relationship, and taking the mean of the adjusted Pearson correlation coefficients as suggested by Falconer and Mackay (1996). Genetic correlations were then calculated using the genetic covariances and the following equation:

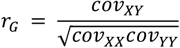

where *cov_XY_* is the genetic ‘cross-covariance’ and *cov_XX_* and *cov_YY_* are the parent-offspring covariances for the individual traits. Standard errors for genetic correlations were calculated using an approximate formula as proposed by Reeve (1955), Robertson (1959) and explained in Falconer and Mackay (1996):

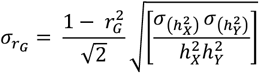

All correlations were estimated using linear regression models that initially included the main effects of temperature and population and an interaction between them, with interaction and population terms removed if they were non-significant. In the majority of cases, population was not significant and this allowed for one correlation value per species.

## Results

Mean trait values differed significantly between thermal environments for each species and generation (*p* = < 0.001). Rearing in a stressful thermal environment resulted in lower fecundity (Supplementary Table 1 and Supplementary Fig. 2), smaller wing size (Supplementary Table 2 and Supplementary Fig. 3), and a rounder, less elongated wing shape when adjusted for size (Supplementary Table 3 and Supplementary Fig. 4) across all species, populations, and generations.

### How does genetic variation change in a stressful thermal environment?

#### Fecundity

Phenotypic variation in fecundity did not differ significantly between thermal environments, but was slightly higher within the stressful environment than the benign environment (Table 1). *CV_A_* estimates could not be calculated for *D. birchii* within the stressful temperature because unexpectedly the offspring did not emerge in this treatment. *CV_A_* and evolvability (*I_A_*) estimates for fecundity were higher than for morphological traits in all instances, and slightly higher in the stressful environment than in the benign environment in *D. serrata* (Table 1 and Fig. 2). Fecundity was found to have a low heritability overall that was slightly higher under the benign than stressful thermal environment in *D. serrata* (Table 1 and Fig. 2).

**Table 1.** Expression of genetic variance parameters for fecundity, wing size, and wing shape; including heritability (*h*^2^), the coefficient of additive variance (*CV_A_*), and evolvability (*I_A_*). Phenotypic (*V_P_*), additive (*V_A_*) and residual (*V_res_*) variances are also shown for the pooled dam and daughter values. Population was not a significant contributor to variance, so one metric was calculated per species from parent-offspring regressions. Bold values indicate a slope significantly different than zero and asterisks indicate significance level after correction for False Discovery Rate (. *P* < 0.1; * *P* < 0.05; *** *P* < 0.001; **** *P* < 0.0001). Parameters could not be calculated for *D. birchii* within the stressful environment because daughters did not develop. *CV_A_* and *I_A_* values shown are x 10^2^. Values for individual relative warp scores for wing shape can be found in Supplementary Table 3.

| Trait | Species | Benign (23°C) |  |  |  |  |  |  | Stressful (28 °C) |  |  |  |  |  |  |
| --- | --- | --- | --- | --- | --- | --- | --- | --- | --- | --- | --- | --- | --- | --- | --- |
| | | $N$ | $h^2 \pm SE$ | $V_P$ | $V_A$ | $V_{res}$ | $CV_A$ | $I_A$ | $N$ | $h^2 \pm SE$ | $V_P$ | $V_A$ | $V_{res}$ | $CV_A$ | $I_A$ |
| <u>Fecundity</u> | *D. birchii* | 86 | $0.148 \pm 0.116$ | 1384.6 | 204.92 | 1179.68 | 15.88 | 2.524 | - | - | - | - | - | - | - |
| | *D. serrata* | 69 | $0.052 \pm 0.124$ | 1105.1 | 57.47 | 1047.64 | 5.26 | 0.276 | 65 | $0.040 \pm 0.139$ | 1879.5 | 75.18 | 1804.32 | 10.17 | 1.032 |
| <u>Wing size</u> | *D. birchii* | 81 | **$0.476 \pm 0.131$** | 0.0005 | 0.0002 | 0.0003 | 0.224 | 0.0005 | - | - | - | - | - | - | - |
| | *D. serrata* | 64 | **$1.000 \pm 0.142^{***}$** | 0.0005 | 0.0005 | 0.0000 | 0.324 | 0.0011 | 62 | $0.226 \pm 0.124$ | 0.0005 | 0.0001 | 0.0004 | 0.156 | 0.0002 |
| <u>Wing shape</u> | *D. birchii* | 81 | **$0.516 \pm 0.22^{****}$** | - | - | - | - | - | - | - | - | - | - | - | - |
|  | *D. serrata* | 64 | **$0.517 \pm 0.25^{***}$** | - | - | - | - | - | 62 | **$0.599 \pm 0.26^*$** | - | - | - | - | - |

**Figure 2:**
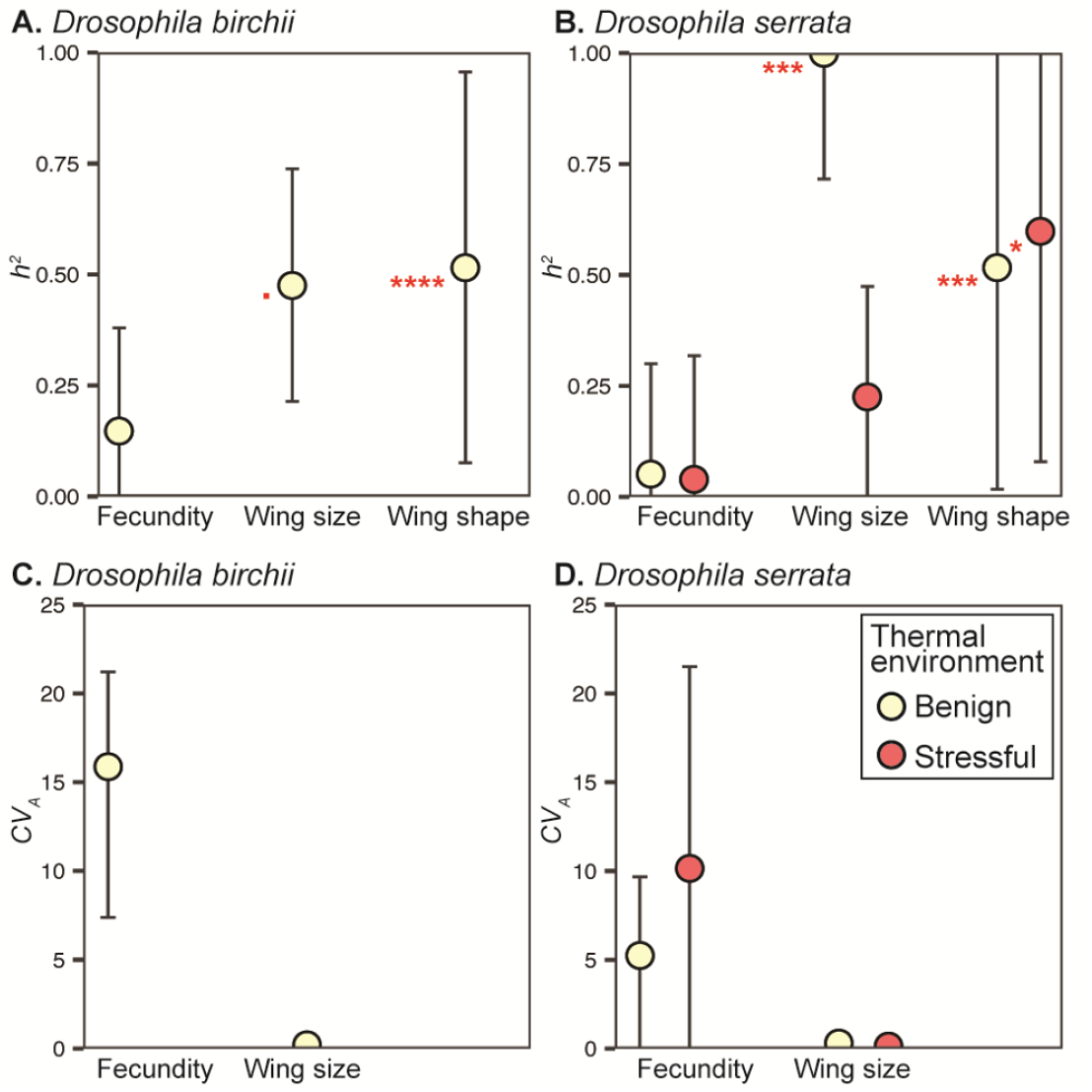
Heritability (*h*^2^) and coefficient of additive variance (*CV_A_*) values for a life history trait and morphological traits across a benign (23°C) and stressful (28°C) thermal environment. Two standardized estimates of additive genetic variance are shown for a life history and two morphological traits in two closely-related species of *Drosophila*. (**A**, **B**) Heritability is standardized by the total genetic variance and (**C**, **D**) coefficient of additive genetic variance is standardized by the trait mean. Evolvability (not shown) will exhibit the same pattern as *CV_A_*. Standard errors (2x) are shown as error-bars, and asterisks indicate significance of the estimate after correction (. *P* < 0.1; * *P* < 0.05; *** *P* < 0.001; **** *P* < 0.0001). Standard errors for *CV_A_* were calculated using the standard error estimates from heritability (see Supplementary Tables 1, 2 for details).

#### Morphological wing traits

Phenotypic variation in wing size differed significantly between thermal environments for dams of both species (*D. serrata: P* = 0.02; *D. birchii: P* = 0.005), and was slightly higher within the benign environment (Supplementary Table 2). Heritability, evolvability, and *CV_A_* estimates were higher within the benign environment than the stressful environment in *D. serrata*, and heritability values were overall much higher for wing size compared to fecundity (Table 1 and Fig. 2).

Phenotypic variation in wing shape variables significantly differed between thermal environments for all RW scores in *D. serrata (P* < 0.005), but did not differ between thermal environments in *D. birchii*. The direction and magnitude of changes in phenotypic variances did not show a consistent pattern across thermal environments (Supplementary Table 3). Wing shape heritability increased within the stressful environment in *D. serrata* (Table 1 and Fig. 2). Heritability in all instances was much higher than for fecundity. In addition, wing size and wing shape evolvabilities and *CV_A_* estimates were all very low compared to fecundity (Fig. 2).

### How does phenotypic correlation of traits change in a stressful environment?

There were no significant phenotypic correlations found between fecundity and wing size after correction for multiple comparisons (Table 2 and Fig. 3).

**Table 2:** Phenotypic correlations between fecundity and wing size. *r_P_* is the phenotypic correlation and the *P*-values were obtained from an *F*-test of the linear regression of one trait on the other and both unadjusted (raw) and adjusted (corrected for using the False Discovery Rate method (Benjamini and Hochberg 1995) are shown. Sample sizes (*N*) indicate the number of individuals used in each correlation. Phenotypic correlations were calculated for each population separately when population was found to be a significant contributor to variation.

| Species | Generation<br>Population | Benign (23°C) |  |  |  | Stressful (28 °C) |  |  |  |
| --- | --- | --- | --- | --- | --- | --- | --- | --- | --- |
| | | $N$ | $r_P$ | $P$-value<br>(raw) | $P$-<br>value | $N$ | $r_P$ | $P$-value<br>(raw) | $P$-<br>value |
| <u>*D. birchii*</u> | Dams | 86 | 0.30 | 0.168 | 0.437 | 78 | 0.22 | 0.346 | 0.647 |
|  | Mt. Lewis | 45 | -0.14 | 0.655 | 0.763 | - | - | - | - |
|  | Paluma | 41 | 0.76 | 0.014* | 0.104 | - | - | - | - |
|  | Daughters | 87 | 0.58 | 0.007** | 0.073 | - | - | - | - |
| <u>*D. serrata*</u> | Dams | 78 | -0.14 | 0.526 | 0.760 | 77 | -0.18 | 0.422 | 0.707 |
|  | Daughters | 67 | 0.12 | 0.628 | 0.763 | 65 | 0.52 | 0.035* | 0.202 |

**Figure 3.**
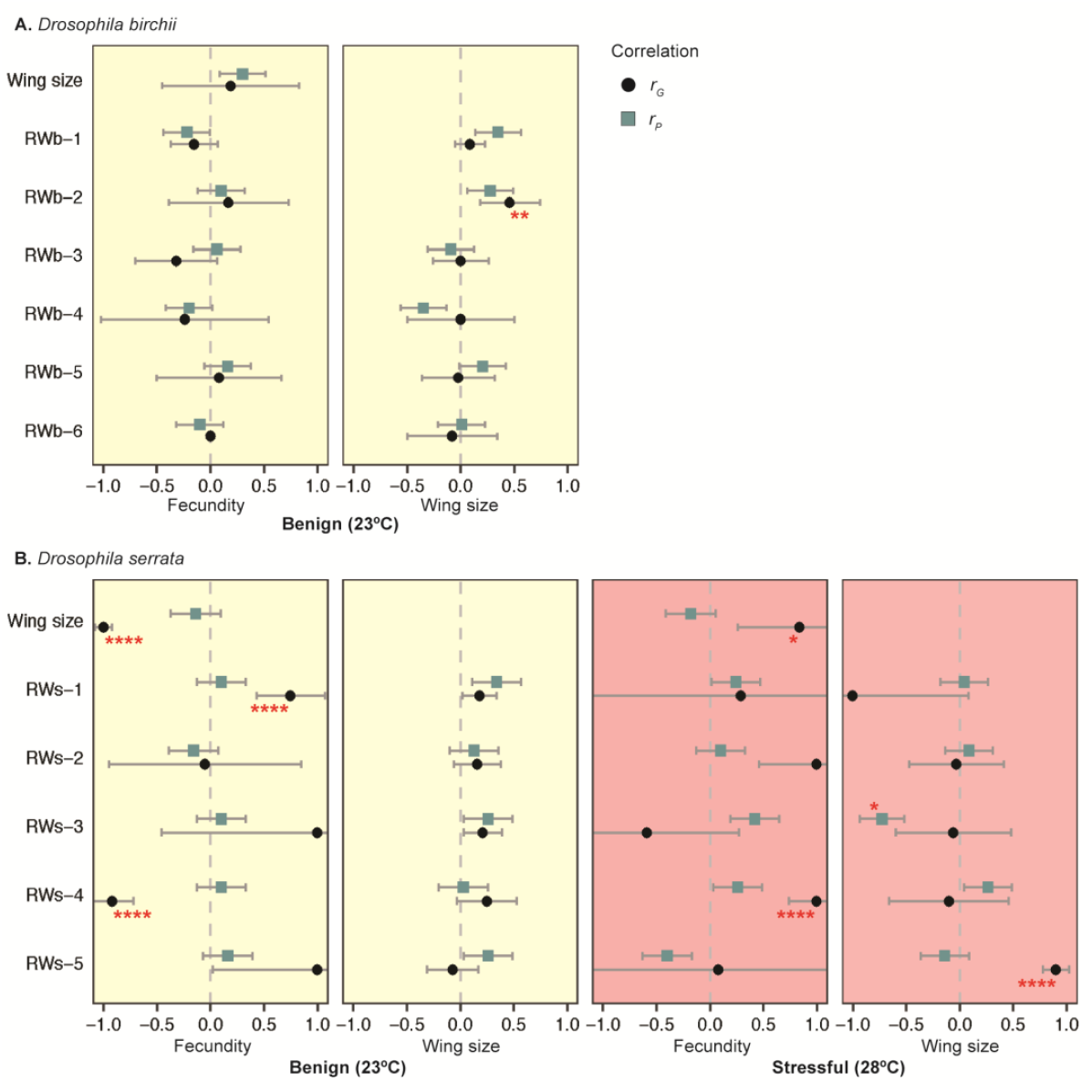
Genetic and phenotypic correlations for fecundity and wing morphology in two sibling-species of *Drosophila*. Genetic correlations (*r_G_*) and phenotypic correlations (*r_P_*) between the trait on the x-axis (fecundity and wing size) and the trait on the y-axis (wing size and wing shape RW scores) across a benign and stressful thermal environment in (**A**) *D. birchii* and (**B**) *D. serrata*. Standard errors (2x) for the correlations are indicated by the grey error bars. Asterisks (in red) denote the correlation is significantly different from 0, obtained from the *z*-statistic calculated from standard errors (Altman and Bland, 2011) and *P*-values have been adjusted by the False Discovery Rate method (Benjamini and Hochberg 1995) (* *P* < 0.05; *** *P* < 0.001; **** *P* < 0.0001). Phenotypic correlations shown are for the dam generation.

Fecundity and wing shape traits exhibited mixed and inconsistent results (Fig. 3 and Supplementary Table 4). There was only one significant phenotypic correlation found between fecundity and a wing shape variable within the daughter generation of the Mt. Lewis population of *D. serrata* under a stressful thermal environment. Allometry was found within the benign environment for the daughter generation of *D. serrata*, indicating wing size and wing shape in this instance are still slightly correlated even after removing effects of size during the GPA analysis (Supplementary Table 4).

### How do genetic covariances and correlations change with environmental stress?

There was no consistent trend detected for genetic covariances and no consistent pattern in genetic correlations (i.e., genetic covariances standardized by individual trait covariances) between species and thermal environments (Fig. 3; Supplementary Tables 5, 6). Genetic correlations in *D. birchii* were generally low (−0.32 < *r_G_* < 0.46), while *D. serrata* traits exhibited high positive and negative genetic correlations, but this was not consistent across environments (Fig. 3).

We did find highly negative and highly positive genetic correlations between fecundity and wing morphometries in *D. serrata* (including values of ± 1.00). However, these often had very wide standard errors and were not always significant. In the benign environment for *D. birchii*, we found a significant positive genetic correlation between wing size and a wing shape variable (RWb-2; *r*_G_ = 0.46 ± 0.14 SE; *P* < 0.01). In the benign environment for *D. serrata*, we found a significant negative genetic correlation between fecundity and wing size (*r*_G_ = −1.00 ± 0.08 SE; *P* < 0.001) and a significant positive (RWs-1; *r*_G_ = 0.75 ± 0.16 SE; *P* < 0.0001) and negative correlation between fecundity and a wing shape variable (RWs-4; *r*_G_ = −0.92 ± 0.10 SE; *P* = 0.0001). In the stressful environment for *D. serrata*, we found a significant positive correlation between fecundity and wing size (*r*_G_ = 0.84 ± 0.29 SE; *P* < 0.05) and fecundity and a wing shape variable (RWs-4; *r*_G_ = 1.00 ± 0.13 SE; *P* < 0.0001; Fig. 3).

## Discussion

The amount of genetic variation in a trait is important for predicting responses of populations and species to climate change as it determines the extent to which a trait can evolve via selection. However, genetic variance and heritability change between environments (Falconer and Mackay 1996; Hoffmann and Schiffer 1998; Hoffmann and Merilä 1999) potentially due to increased environmental variance and reduced additive genetic variance, genotype-by-environment interactions that affect cross-environment genetic correlations, and cryptic genetic variation (Fischer et al. 2020). It is essential to recognise and incorporate this consideration into climate change adaptation research (Shaw 2019), but consistent and predictable patterns have not been detected. It is unclear whether such patterns exist or whether genetic variance and heritability must always be considered in the context of specific traits, populations and environments. Here, we show that temperature stress can alter the heritability, coefficient of additive genetic variation, and evolvability, of both fecundity and morphological traits in two closely-related species of *Drosophila* (one being a generalist and one being a specialist). However, we found no consistent pattern in the direction of change in additive genetic variance and phenotypic and genetic covariances across thermal environments.

First, we confirmed that the warmer (‘stressful’) thermal environment did indeed induce stress in both species, as demonstrated by lower fecundity, smaller size, and a significantly different wing shape (Supplementary Tables 1–3 and Supplementary Figs. 2–4). In addition, the specialist species (*D. birchii*) failed to develop offspring within the stressful thermal environment. Although there were no experimental differences between the dam and daughter generation that might have caused this, it is possible that maternal effects induced by development within a stressful environment prevented the production of viable offspring. This could alternatively be a paternal effect as it has been shown that *D. birchii* sperm is very sensitive to thermal stress during development (Saxon et al. 2018), and because we attempted to control for maternal effects by developing the initial parental generation (i.e., P in Fig. 1) in the same thermal environment as dams and daughters. Although not a direct aim of this paper, measuring the viability of offspring within a stressful environment is relevant to many evolutionary studies (both in the laboratory and in the field). This is because many studies estimate fitness by measuring the number of offspring directly, but the viability of those offspring are what will maintain the long-term fitness of a population.

Second, we found lower heritability in the stressful compared to the benign thermal environment for fecundity and wing size, although not in wing shape in *D. serrata* (Table 1; Fig. 2B). This corroborates a large number of previous studies that show heritability declined under stressful conditions (e.g., Hoffmann and Schiffer 1998; Kristensen et al. 2015; and reviewed in Hoffmann and Parsons 1991; Hoffmann and Merilä 1999; Charmantier and Garant 2005; Rowiński and Rogell 2017). This has important implications for species living close to their upper thermal limits (like many species in the tropics (Deutsch et al. 2008; Kingsolver et al. 2013) because even a small change in environmental conditions may induce a large amount of stress, and these results suggest adaptive potential is reduced under stressful temperatures.

However, heritability has been shown to have inherent issues when comparing between environments, as non-additive genetic and environmental variation contribute to it (Houle 1992). To address this problem, we also investigated the coefficient of additive genetic variation (*CV_A_*) and evolvability (*I_A_*). These are often more appropriate estimates to use when comparing genetic variation and evolvability across traits and environments, as they are not affected by non-additive sources of environmental variance (Houle 1992; Bubliy and Loeschcke 2002; Garcia-Gonzalez et al. 2012). Specifically, while heritability tells us the expected absolute change in a trait mean under a given strength of selection from one generation to the next, evolvability predicts the relative change as a percentage of the trait mean (Hansen et al. 2003; Hansen et al. 2011; Garcia-Gonzalez et al. 2012). *CV_A_* and *I_A_* were higher under the stressful environment for fecundity, while the opposite was true for wing size (Fig. 2D). Therefore, while the heritability values suggest that the response to selection on fecundity and wing size will decrease under stressful temperatures, *CV_A_* and *I_A_* suggest that fecundity has greater relative evolutionary potential under the stressful environment than the benign environment and the opposite is true for wing size (Fig. 2). Although it seems heritability and *CV_A_* values may be contradictory, it could be that while the absolute change in fitness (i.e., heritability) will be less in the stressful environment for fecundity, there will be a greater relative increase in fitness in the stressful environment because mean fitness is lower — but this could result in a smaller absolute change in trait mean thus corroborating the heritability results. However, *CV_A_* and *I_A_* values are still important metrics to consider because they will always change in the same direction as *V_A_*. Alternatively, heritability does not necessarily change in the same direction as *V_A_* because effects that increase *V_A_* often increase total variance, which in turn will decrease heritability.

An increase in additive genetic variance under stressful temperatures for the measured fitness trait (i.e., fecundity) is advantageous for these *Drosophila* species, both of which live near critical thermal limits (Kellermann et al. 2009; Overgaard et al., 2011). Interestingly, these results are consistent with a recent meta-analysis (Rowiński and Rogell 2017), which showed that the coefficient of genetic variance (*CV_A_*) was higher under stressful conditions for life history traits but not for morphological traits. In terms of wing size, it should be noted that the measured *CV_A_* differed from values previously measured in *D. birchii* (Kellermann et al. 2006), where *CV_A_* was relatively two-fold higher than what we found here. However, in the previous experiment (Kellermann et al. 2006), wing size was measured from flies reared in a benign environment at 25°C (compared to the benign environment measured in this study at 23°C), potentially indicating that even a slight difference in thermal environments can affect estimates of genetic variance. The reported *V_A_* and *V_P_* values indicate differences in *CV_A_* between this study and Kellermann et al. (2006) are due to an increased *V_A_* in their study and not a difference in trait mean that could also induce larger *CV_A_* values (if the trait mean was lower). Collectively, this, along with the other results discussed here, reveal that environmental interactions (that are included in estimating heritability but not *CV_A_* and *I_A_*), potentially play a very large role in shaping the amount of additive genetic variance that selection can act upon.

Overall, our heritability values are similar to those reported for fecundity, wing size, and wing shape for *Drosophila* (Gilchrist and Partridge 1999; Hoffmann and Shirriffs 2002; Moraes et al. 2004; Kellermann et al. 2006). Additionally, we examined the differences in genetic variation between fecundity and morphological traits, since patterns in heritability and additive variance (*CV_A_* and *I_A_*) were contradictory. We found that heritabilities were higher for the wing morphology traits than for fecundity (Fig. 2A, B). This coincides with the majority of literature that show morphological traits often have higher heritabilities than life history traits (Mousseau and Roff 1987; but for opposing example see Sgrò and Hoffmann 1998c). In direct contrast to this, *CV_A_* and *I_A_* were both magnitudes larger for fecundity than what was found for wing morphology (Fig. 2C, D). This finding supports theory proposed by Houle (1992); that life history traits may have a higher evolvability than morphological traits. Under the benign environment, *CV_A_* and *I_A_* for fecundity were more than 94% higher than for wing size in both species, and in the stressful environment, fecundity exhibited a *CV_A_* and *I_A_* that was approximately 80% higher than for wing size for *D. serrata* (Fig. 2).

The low heritability values detected for fecundity are consistent with classic theory that suggests ultimate fitness traits will exhibit low heritabilities due to directional selection that fixes beneficial alleles and erodes additive and residual variance (Mousseau and Roff 1987; Falconer and Mackay 1996; Merilä and Sheldon 1999). However, in direct contrast to this, we found that additive variance was actually significantly higher in fecundity where *h^2^* was low. When examining residual variance (*V_res_* = *V_P_* – *V_A_*), it becomes evident that increased residual variance is responsible for a reduced heritability in fecundity, rather than eroded additive genetic variance (Table 1; Kruuk et al. 2000; Merilä and Sheldon 2000; McCleery et al. 2004; Moraes et al. 2004). In a study examining how residual and additive variance contributes to heritability values across fitness and morphological traits, Merilä & Sheldon (2000) found fitness traits generally exhibit a higher residual variance compared to morphological traits due to an accumulation of non-additive genetic and early environmental effects. These results support their findings and emphasize the importance of considering trait type when examining how selection shapes additive genetic variance.

An additional aim of this study was to determine whether an easy-to-measure morphological trait can be used as a proxy for fecundity across environments. To examine this, we looked at phenotypic and genetic correlations between fecundity and wing morphology. Although it has been shown that wing length correlates with fecundity in benign environments (Chiang and Hodson 1950; Tantawy and Vetukhiv 1960; Santos et al. 1992; Woods et al. 2002), recent studies have found both evidence for (Woods et al. 2002) or a lack of evidence for (Sgrò and Hoffmann 1998a) positive relationships between wing length, wing width, and fecundity in stressful environments. Here, unadjusted significance tests are suggestive of significant phenotypic correlations between fecundity and wing size in the benign environment for one population of *D. birchii* dams and for *D. birchii* daughters; and in the stressful thermal environment for *D. serrata* daughters. However, these became insignificant after we corrected for False Discovery Rate (Table 2). Although this is a conservative method for multiple comparison in terms of type II errors, the results suggest we cannot use wing size as a proxy for fecundity for these populations in these thermal environments (Fig. 3).

Genetic correlations were all fairly low in *D. birchii*, but highly-positive and highly-negative correlations were found in both environments for *D. serrata* (Fig. 3; Supplementary Table 6). Most interestingly in *D. serrata*, fecundity and wing size were significantly negatively-correlated in the benign environment and significantly positively-correlated in the stressful environment. A significant genetic correlation between a pair of traits suggests that the traits are genetically associated through linkage or pleiotropy (influenced by a common locus or loci; Wilson et al. 2010). However, a change in the magnitude or sign of genetic correlations across environments suggests that this genetic association is environment-specific (Falconer and Mackay 1996; see Gutteling et al. 2007 for example). So, while a positive correlation between fecundity and wing size in the stressful environment may indicate that the same gene underlies both traits or the genes influencing both traits are in linkage disequilibrium (Wood and Brodie 2016); a negative correlation in the benign environment may indicate antagonistic pleiotropy between them if this data was looked at independently. However, the drastic change between thermal environments suggests there are environment-specific gene effects that affect these correlations.

Additionally, when examining phenotypic correlations and genetic correlations together, we did not find phenotypic correlations that were similar to significant genetic correlations (Fig. 3). This suggests that the environment may be masking phenotypic correlations. The large standard errors associated with many of the genetic correlations also suggest that we may lack sufficient power to detect genetic correlations in some cases. Very large sample sizes are needed in quantitative genetic experiments to estimate heritabilities and genetic correlations with a high degree of precision (Roff 1995; Falconer and Mackay 1996). This is hard to achieve due to logistical challenges, and may partly explain why there is much variation across species, populations, and traits in the literature.

### Conclusion

Here, we found that genetic variance and phenotypic and genetic correlations change across thermal environments. However, the direction of these changes was not always consistent across traits, closely-related species, populations within a species, or even generations. This suggests that researchers need to examine adaptive potential specific to their environment, species, and populations if they hope to obtain accurate parameters to predict evolutionary potential. The type of data collected here should represent a starting point for researchers aiming to do so.

Additionally, researchers need to be aware that high genetic variation does not necessarily indicate an increased evolutionary response. Although it is assumed that selection has a strong effect when genetic variation is high and a weak effect when genetic variation is low (when all other factors remain the same), there has been limited evidence showing how selection and genetic variance interact and the studies that have looked at their relationship report a fairly weak association (Wood and Brodie 2016; Ramakers et al. 2018). Future research needs to consider how evolutionary potential is affected by the environment. We show here that genetic variance is highly dependent on temperature and it is accepted that selection is directly mediated by the environment. Yet, specifically in terms of stressful temperatures, a meta-analysis on how selection and genetic variance are coupled found temperature is likely to affect the amount of genetic variation in a population more than the strength of selection (Wood and Brodie 2016). Wood and Brodie (2016) found that temperature affected the amount of genetic variation and the strength of selection in both morphological and fitness traits asymmetrically; meaning the measured impact of temperature stress on genetic variation does not necessarily predict the magnitude of the evolutionary change. Researchers should examine how both genetic variance and selective force (both strength and directionality of selection) is influenced by specific environments to determine the adaptive potential of species to climate change. If a highly positive correlation exists between the two, environmental change would increase both, directly causing increased adaptation; and predictions on how species will adapt to changing environments would be more straightforward (Wood and Brodie 2016; Ramakers et al. 2018; Fischer et al. 2020). Genetic correlations also need to be considered in this context. A negative genetic correlation between two traits will constrain evolution on one trait even with an increase in genetic variation and a positive selection differential (and vice versa; Conner 2012; Wood and Brodie 2016). An additional consideration is that the underlying genetic architecture of the trait (polygenic or large-effect loci) should be considered. For example, polygenic traits have been shown to produce greater long-term population viability than in traits affected by large-effect alleles when heritability and the selective force is constant (e.g., Kardos and Luikart 2021). Generally, life-history traits are thought to be polygenic in comparison to large-effect phenotypic traits related to morphology, indicating another reason why trait type needs to be considered when investigating adaptive potential.

In conclusion, although we present clear evidence that stressful temperatures affect genetic variation, we did not detect a consistent pattern to that change. These results suggest that adaptive potential cannot be generalized across environments, closely-related species or populations and needs to be considered on a case-by-case basis, specific to the trait in question, and by using a multivariate approach.

## Acknowledgements

We would like to thank Malaika Chawla, Nicholas Bail, Naomi Laven, Natalie Swinhoe, and Paula Strickland for assistance in data collection, specifically for helping with the task of egg-counting. We thank Jodie Betts for technical assistance. This study was supported by grants from the American Society of Naturalists, the Ecological Society of Australia, the Wet Tropics Management Authority, James Cook University, and the American Australian Association to JC, as well as an Australian Research Council Discovery Grant (DE130100218) to MH.

## Conflict of Interest

The authors declare there are no conflicts of interest.

## Data archiving

Data will be archived in the Dryad data repository if paper is accepted for publication.

## Notes

### Competing Interest Statement

The authors have declared no competing interest.

