## Supplementary Material for "The predictive potential of key adaptation parameters and proxy fitness traits between benign and stressful thermal environments"

### List of supplementary tables

**Supplementary Table 1**: Sample sizes, mean trait values, phenotypic variances ($V_P$), additive variances $V_A$), residual variances ($V_{res}$), coefficient of additive variance ($CV_A$), and evolvabilities ($I_A$) for fecundity .. 2

**Supplementary Table 1. Sample sizes, mean trait values, phenotypic variances ($V_P$), additive variances ($V_A$), residual variances ($V_{res}$), coefficient of additive variance ($CV_A$), and evolvabilities ($I_A$) for fitness as measured by fecundity in a benign (23ºC) and stressful (28ºC) environment**.

Standard errors for $CV_A$ were calculated by adding and subtracting the SE of heritability from the heritability value and then calculated $V_A$ from each estimate (i.e., $V_A = h^2 \times V_P$) to obtain an approximate lower standard error (*LSE*) and upper standard error (*USE*).

| | | | | | 23°C (Benign environment) | | | | |
|---|---|---|---|---|---|---|---|---|---|
| Species | Generation | *N* | *mean* ± *SE* (no. of offspring) | $V_P$ | $V_A$ | $V_{res}$ | $CV_A$ x $10^2$ | *SE* $CV_A$ *x 10 [LSE; USE]* | $I_A$ x $10^2$ |
| *D. birchii* | Pooled | 179 | 90.1 ± 2.78 | 1384.6 | 204.921 | 1179.679 | 15.888 | [7.39; 21.22] | 2.524 |
| | Dams | 91 | 101.5 ± 2.60 | 630.4 | - | - | - | - | - |
| | Daughters | 88 | 78.4 ± 4.70 | 1906.3 | - | - | - | - | - |
| *D. serrata* | Pooled | 154 | 144.19 ± 2.67 | 1105.1 | 57.465 | 1047.635 | 5.257 | [0; 9.67] | 0.276 |
| | Dams | 81 | 131.2 ± 3.50 | 991.4 | - | - | - | - | - |
| | Daughters | 73 | 158.7 ± 3.40 | 843.4 | - | - | - | - | - |
| | | | | | 28°C (Stressful environment) | | | | |
| *D. birchii* | Pooled | - | - | - | - | - | - | - | - |
| | Dams | 86 | 16.20 ± 1.86 | 299.8 | - | - | - | - | - |
| | Daughters | - | - | - | - | - | - | - | - |
| *D. serrata* | Pooled | 152 | 85.34 ± 3.51 | 1879.5 | 75.180 | 1804.320 | 10.165 | [0; 21.50] | 1.032 |
| | Dams | 84 | 76.8 ± 4.30 | 1548.1 | - | - | - | - | - |
| | Daughters | 68 | 95.9 ± 5.60 | 2111.7 | - | - | - | - | - |

**Supplementary Table 2. Sample sizes, mean trait values, phenotypic variances ($V_P$), additive variances ($V_A$), residual variances ($V_{res}$), coefficient of additive variance ($CV_A$), and evolvabilities ($I_A$) for wing size in a benign (23ºC) and stressful (28ºC) environment**.

Wing size is shown as the log centroid size, which produces an arbitrary unit of measurement for comparison purposes. Standard errors for $CV_A$ were calculated by adding and subtracting the SE of heritability from the heritability value and then calculated $V_A$ from each estimate (i.e., $V_A = h^2 \times V_P$) to obtain an approximate lower standard error ($LSE$) and upper standard error ($USE$).

| Species | Generation | *N* | *mean ± SE* (log centroid size) | $V_P$ | $V_A$ | $V_{res}$ | $CV_A \times 10^2$ | *SE* $CV_A$ *x 10 [LSE; USE]* | $I_A \times 10^2$ |
|---|---|---|---|---|---|---|---|---|---|
| | | | 23°C (Benign environment) | | | | | | |
| D. birchii | Pooled | 173 | 6.8786 ± 0.0016 | 0.0005 | 0.0002 | 0.0003 | 0.224 | [0.191; 0.253] | 0.001 |
| | Dams | 86 | 6.8813 ± 0.0026 | 0.0006 | - | - | - | - | - |
| | Daughters | 87 | 6.8758 ± 0.0020 | 0.0004 | - | - | - | - | - |
| D. serrata | Pooled | 145 | 6.9076 ± 0.0020 | 0.0005 | 0.0005 | 0.0000 | 0.324 | [0.299; 0.346] | 0.001 |
| | Dams | 79 | 6.9065 ± 0.0024 | 0.0004 | - | - | - | - | - |
| | Daughters | 66 | 6.9088 ± 0.0034 | 0.0007 | - | - | - | - | - |
| | | | 28°C (Stressful environment) | | | | | | |
| D. birchii | Pooled | - | - | - | - | - | - | - | - |
| | Dams | 78 | 6.7723 ± 0.0026 | 0.0005 | - | - | - | - | - |
| | Daughters | - | - | - | - | - | - | - | - |
| D. serrata | Pooled | 147 | 6.8159 ± 0.0019 | 0.0005 | 0.0001 | 0.0004 | 0.156 | [0.105; 0.194] | 0.0002 |
| | Dams | 81 | 6.8091 ± 0.0026 | 0.0005 | - | - | - | - | - |
| | Daughters | 66 | 6.8244 ± 0.0025 | 0.0004 | - | - | - | - | - |

**Supplementary Table 3. Sample sizes, mean trait values, heritabilities ($h^2$), phenotypic variances ($V_P$), additive variances ($V_A$), and residual variances ($V_{res}$) for the relative warp (RW) parameters for wing shape in a benign (23ºC) and stressful (28ºC) environment**.

Heritabilities shown in bold are significantly different from zero and the asterisks indicate the significance level for adjusted $P$-values (adjusted by False Discovery Rate; $*P < 0.05$, $**P < 0.01$, $***P < 0.001$, $****P < 0.0001$). The percentage of variation that each RW score accounts for is also shown.

| | | | | 23°C (Benign environment) | | | | | |
|---|---|---|---|---|---|---|---|---|---|
| Species | | Generation | *N* | *mean ± SE* | $h^2$ | $V_P$ | $V_A$ | $V_{res}$ | % variation |
| <u>*D. birchii*</u> | RWb-1 | Pooled | 173 | 0.0061 ± 0.0008 | **0.412** ± 0.22* | 1.12E-04 | 4.66E-05 | 6.58E-05 | 34.47 |
| | | Dams | 86 | 0.0069 ± 0.0011 | - | 1.12E-04 | - | - | - |
| | | Daughters | 87 | 0.0053 ±0.0010 | - | 8.94E-05 | - | - | - |
| | RWb-2 | Pooled | 173 | 0.0005 ± 0.0007 | **0.356** ± 0.22** | 1.12E-04 | 3.05E-05 | 8.12E-05 | 16.97 |
| | | Dams | 86 | 0.0017 ± 0.0011 | - | 7.21E-05 | - | - | - |
| | | Daughters | 87 | -0.0008 ± 0.0008 | - | 5.78E-05 | - | - | - |
| | RWb-3 | Pooled | 173 | -0.001 ± 0.0007 | **0.680** ± 0.22*** | 7.21E-05 | 4.90E-05 | 2.31E-05 | 13.09 |
| | | Dams | 86 | -0.009 ± 0.0010 | - | 7.21E-05 | - | - | - |
| | | Daughters | 87 | -0.0011 ±0.0008 | - | 5.33E-05 | - | - | - |
| | RWb-4 | Pooled | 164 | 0.0007 ± 0.0005 | 0.142 ± 0.22 | 3.73E-05 | 5.30E-06 | 3.20E-05 | 6.54 |
| | | Dams | 86 | 0.0011 ± 0.0007 | - | 3.73E-05 | - | - | - |
| | | Daughters | 87 | 0.0003 ± 0.0006 | - | 3.03E-05 | - | - | - |
| | RWb-5 | Pooled | 173 | -0.0002 ± 0.0004 | **0.457** ± 0.22* | 3.31E-05 | 1.51E-05 | 1.80E-05 | 6.07 |
| | | Dams | 86 | -0.00004 ± 0.0006 | - | 3.58E-05 | - | - | - |
| | | Daughters | 87 | -0.0003 ± 0.0006 | - | 3.07E-05 | - | - | - |
| | RWb-6 | Pooled | 173 | 0.0003 ± 0.0004 | 0.035 ± 0.22 | 2.62E-05 | 6.89E-10 | 2.62E-05 | 5.55 |
| | | Dams | 86 | 0.0005 ± 0.0006 | - | 2.75E-05 | - | - | - |
| | | Daughters | 87 | 0.00003 ± 0.0005 | - | 2.52E-05 | - | - | - |
| <u>*D. serrata*</u> | RWs-1 | Pooled | 145 | 0.0116 ± 0.0009 | **1.00** ± 0.25**** | 1.05E-04 | 1.05E-04 | 0.00E+00 | 41.11 |
| | | Dams | 79 | 0.010 ± 0.0010 | - | 1.04E-04 | - | - | - |
| | | Daughters | 66 | 0.013 ± 0.0010 | - | 1.02E-04 | - | - | - |
| | RWs-2 | Pooled | 145 | 0.0002 ± 0.0007 | **0.836** ± 0.25* | 7.56E-05 | 6.32E-05 | 1.24E-05 | 15.96 |
| | | Dams | 79 | 0.0002 ± 0.0090 | - | 6.63E-05 | - | - | - |
| | | Daughters | 66 | 0.0003 ± 0.0110 | - | 8.77E-05 | - | - | - |
| | RWs-3 | Pooled | 145 | -0.0017 ± 0.0006 | **1.00** ± 0.25**** | 5.79E-05 | 5.79E-05 | 0.00E+00 | 10.27 |
| | | Dams | 79 | -0.0025 ± 0.0008 | - | 5.62E-05 | - | - | - |
| | | Daughters | 66 | -0.008 ± 0.0009 | - | 5.90E-05 | - | - | - |
| | RWs-4 | Pooled | 145 | -0.0004 ± 0.0005 | 0.514 ± 0.25 | 4.01E-05 | 2.06E-05 | 1.95E-05 | 7.24 |
| | | Dams | 79 | -0.0011 ± 0.0006 | - | 3.00E-05 | - | - | - |
| | | Daughters | 66 | 0.0005 ± 0.0009 | - | 5.12E-05 | - | - | - |
| | RWs-5 | Pooled | 145 | 0.0008 ± 0.0005 | 0.510 ± 0.25 | 3.58E-05 | 1.82E-05 | 1.75E-05 | 6.45 |
| | | Dams | 79 | 0.0008 ± 0.0007 | - | 3.96E-05 | - | - | - |
| | | Daughters | 66 | 0.0008 ± 0.0005 | - | 3.58E-05 | - | - | - |

| Species | | Generation | *N* | *mean ± SE* | $h^2$ | $V_P$ | $V_A$ | $V_{res}$ | % variation |
|---|---|---|---|---|---|---|---|---|---|
| 28°C (Stressful environment) | | | | | | | | | |
| *D. birchii* | RWb-1 | Pooled | - | - | - | - | - | - | 34.47 |
| | | Dams | 78 | -0.0135 ± 0.0011 | - | 9.41E-05 | - | - | - |
| | | Daughters | - | - | - | - | - | - | - |
| | RWb-2 | Pooled | - | - | - | - | - | - | 16.97 |
| | | Dams | 78 | -0.001 ± 0.0011 | - | 9.63E-05 | - | - | - |
| | | Daughters | - | - | - | - | - | - | - |
| | RWb-3 | Pooled | - | - | - | - | - | - | 13.09 |
| | | Dams | 78 | 0.0022 ± 0.0008 | - | 5.50E-05 | - | - | - |
| | | Daughters | - | - | - | - | - | - | - |
| | RWb-4 | Pooled | - | - | - | - | - | - | 6.54 |
| | | Dams | 78 | -0.0016 ± 0.0006 | - | 2.45E-05 | - | - | - |
| | | Daughters | - | - | - | - | - | - | - |
| | RWb-5 | Pooled | - | - | - | - | - | - | 6.07 |
| | | Dams | 78 | 0.0004 ± 0.0006 | - | 2.93E-05 | - | - | - |
| | | Daughters | - | - | - | - | - | - | - |
| | RWb-6 | Pooled | - | - | - | - | - | - | 5.55 |
| | | Dams | 78 | -0.0006 ± 0.0007 | - | 3.55E-05 | - | - | - |
| | | Daughters | - | - | - | - | - | - | - |
| *D. serrata* | RWs-1 | Pooled | 147 | -0.0116 ± 0.0009 | **0.268** ±0.25** | 1.06E-04 | 2.84E-05 | 7.76E-05 | 41.11 |
| | | Dams | 81 | -0.015 ± 0.001 | - | 1.14E-04 | - | - | - |
| | | Daughters | 66 | -0.008 ± 0.001 | - | 7.11E-05 | - | - | - |
| | RWs-2 | Pooled | 147 | -0.0002 ± 0.0009 | **0.668** ±0.25* | 1.12E-04 | 7.45E-05 | 3.70E-05 | 15.96 |
| | | Dams | 81 | -0.0015 ± 0.0012 | - | 1.19E-04 | - | - | - |
| | | Daughters | 66 | 0.0014 ± 0.0012 | - | 1.00E-04 | - | - | - |
| | RWs-3 | Pooled | 147 | 0.0017 ± 0.0006 | 0.576 ±0.25 | 5.67E-05 | 3.27E-05 | 2.40E-05 | 10.27 |
| | | Dams | 81 | 0.0024 ± 0.0008 | - | 4.82E-05 | - | - | - |
| | | Daughters | 66 | 0.0009 ± 0.0010 | - | 6.69E-05 | - | - | - |
| | RWs-4 | Pooled | 147 | 0.0004 ± 0.0006 | 0.414 ±0.25 | 4.45E-05 | 1.84E-05 | 2.61E-05 | 7.24 |
| | | Dams | 81 | 0.00001 ± 0.0008 | - | 5.00E-05 | - | - | - |
| | | Daughters | 66 | 0.0008 ± 0.0008 | - | 3.81E-05 | - | - | - |
| | RWs-5 | Pooled | 147 | -0.0008 ± 0.0005 | 0.430 ±0.25 | 3.86E-05 | 1.66E-05 | 2.20E-05 | 6.45 |
| | | Dams | 81 | 0.00001 ± 0.0007 | - | 3.91E-05 | - | - | - |
| | | Daughters | 66 | -0.0018 ± 0.0007 | - | 3.66E-05 | - | - | - |

**Supplementary Table 4: Phenotypic correlations between traits in *D. birchii* and *D. serrata* reared under two temperatures.**

$r_P$ is the phenotypic correlation and the *P*-value is adjusted by False Discovery Rate and was obtained from a *F*-test of the linear regression of one trait on the other. Bold values indicate significance and significance level is shown in asterisks (**P* < 0.05, ***P* < 0.01, ****P* < 0.001, *****P* < 0.0001). In addition, a *F*-test was conducted to determine if the interaction of population with this regression was significant, which tests whether the phenotypic correlation varies between the two populations of a species. Where there was a significant interaction, $r_P$ was estimated for each population individually.

| Trait 1 ~ Trait 2 | Species | Trait | Generation Population | $r_P$ | SE | P-value | $r_P$ | SE | P-value |
|---|---|---|---|---|---|---|---|---|---|
| | | | | Benign (23°C) | | | Stressful (28°C) | | |
| Fecundity ~ Wing size | D. birchii | | Dams | 0.30 | 0.108 | 0.437 | 0.22 | 0.117 | 0.647 |
| | | | Mt. Lewis | -0.14 | 0.147 | 0.763 | - | - | - |
| | | | Paluma | 0.76 | 0.147 | 0.104 | - | - | - |
| | | | Daughters | 0.58 | 0.103 | 0.073 | - | - | - |
| | D. serrata | | Dams | -0.14 | 0.117 | 0.760 | -0.18 | 0.117 | 0.707 |
| | | | Daughters | 0.12 | 0.123 | 0.763 | 0.52 | 0.125 | 0.202 |
| Fecundity ~ Wing shape | D. birchii | RWb-1 | Dams | -0.22 | 0.108 | 0.625 | - | - | - |
| | | | Daughters | 0.10 | 0.108 | 0.763 | - | - | - |
| | | RWb-2 | Dams | 0.10 | 0.109 | 0.763 | - | - | - |
| | | | Daughters | 0.20 | 0.108 | 0.647 | - | - | - |
| | | RWb-3 | Dams | 0.06 | 0.109 | 0.834 | - | - | - |
| | | | Daughters | -0.26 | 0.108 | 0.507 | - | - | - |
| | | RWb-4 | Dams | -0.20 | 0.109 | 0.647 | - | - | - |
| | | | Mt Lewis | -0.76 | - | 0.104 | - | - | - |
| | | | Paluma | 0.04 | - | 0.507 | - | - | - |
| | | | Daughters | -0.16 | 0.108 | 0.707 | - | - | - |
| | | RWb-5 | Dams | 0.16 | 0.109 | 0.707 | - | - | - |
| | | | Daughters | -0.40 | 0.106 | 0.243 | - | - | - |
| | | RWb-6 | Dams | -0.10 | 0.109 | 0.763 | - | - | - |
| | | | Daughters | 0.42 | 0.106 | 0.218 | - | - | - |
| | D. serrata | RWs-1 | Dams | 0.10 | 0.115 | 0.763 | 0.24 | 0.11 | 0.625 |
| | | | Daughters | 0.08 | 0.126 | 0.837 | -0.22 | 0.11 | 0.647 |
| | | RWs-2 | Dams | -0.16 | 0.115 | 0.707 | 0.10 | 0.11 | 0.787 |
| | | | Daughters | 0.04 | 0.126 | 0.902 | -0.52 | 0.12 | 0.203 |
| | | RWs-3 | Dams | 0.10 | 0.115 | 0.763 | 0.42 | 0.11 | 0.247 |
| | | | Daughters | -0.04 | 0.125 | 0.902 | 0.06 | 0.13 | 0.856 |
| | | RWs-4 | Dams | 0.10 | 0.115 | 0.763 | 0.26 | 0.11 | 0.581 |
| | | | Daughters | -0.14 | 0.125 | 0.763 | 0.54 | 0.12 | 0.202 |
| | | RWs-5 | Dams | 0.16 | 0.115 | 0.760 | -0.40 | 0.11 | 0.283 |
| | | | Daughters | 0.36 | 0.125 | 0.437 | 0.48 | 0.12 | 0.218 |
| | | | Mt Lewis | - | - | - | **0.98*** | - | 0.049 |
| | | | Paluma | - | - | - | -0.06 | - | 0.897 |
| Wing size ~ Wing shape | D. birchii | | Dams | 0.26 | - | 0.437 | 0.33 | - | 0.218 |
| | | | Daughters | 0.39 | - | 0.065 | - | - | - |
| | D. serrata | | Dams | 0.43 | - | 0.052 | 0.28 | - | 0.401 |
| | | | Daughters | **0.59*** | - | 0.049 | 0.28 | - | 0.565 |

**Supplementary Table 5: Genetic covariances and correlations between traits in *D. birchii* and *D. serrata* reared under a benign (23ºC) and stressful (28 ºC) environment.**

Genetic covariances and correlations were calculated in both directions, meaning one trait in the dam was regressed on the other trait in the daughters and vice versa. *N* indicates the number of family pairs used in the regression, the slope for the regression is indicated by $\beta$, the genetic covariance of one trait on the other is $cov_{XY}$, the covariances for the individual traits are shown as $cov_{XX}$ and $cov_{YY}$, and the genetic correlation ($r_G$) was calculated using the equation set forth in Falconer and Mackay (1996). The adjusted *P*-value for the regression is noted (adjusted by False Discovery Rate), as well as the *P*-value for a *F*-test on the interaction between population and the trait value (*P*-value$_{pop}$). If population was significant, individual parameters were estimated for each. Genetic correlations shown in the paper were calculated from the mean of the genetic covariances in both directions.

| Species | Dam - daughter Population | Benign (23ºC) | | | | | | | |
|---|---|---|---|---|---|---|---|---|---|
| | | *N* | $\beta$ | $cov_{XY}$ | $cov_{XX}$ | $cov_{YY}$ | $r_G$ | *P*-value | *P*-value$_{pop}$ |
| D. birchii | Fecundity–Wing size | 85 | 0.14 | 0.280 | 0.148 | 0.476 | 1.05 | 0.869 | 0.863 |
| | Wing size–Fecundity | 82 | -0.09 | -0.180 | 0.476 | 0.148 | -0.68 | 0.869 | 0.335 |
| | Fecundity–RWb-1 | 85 | -0.07 | -0.140 | 0.148 | 0.462 | -0.54 | 0.869 | 0.407 |
| | RWb-1–Fecundity | 82 | 0.03 | 0.060 | 0.462 | 0.148 | 0.23 | 0.932 | 0.282 |
| | Fecundity–RWb-2 | 85 | 0.07 | 0.140 | 0.148 | 0.356 | 0.61 | 0.869 | 0.354 |
| | RWb-2–Fecundity | 82 | -0.03 | -0.060 | 0.356 | 0.148 | -0.26 | 0.932 | 0.671 |
| | Fecundity–RWb-3 | 85 | -0.13 | -0.260 | 0.148 | 0.680 | -0.82 | 0.869 | 0.867 |
| | RWb-3–Fecundity | 82 | 0.03 | 0.060 | 0.680 | 0.148 | 0.19 | 0.932 | 0.485 |
| | Fecundity–RWb-4 | 85 | -0.07 | -0.140 | 0.148 | 0.284 | -0.68 | 0.869 | 0.141 |
| | RWb-4–Fecundity | 82 | 0.02 | 0.040 | 0.284 | 0.148 | 0.20 | 0.960 | 0.273 |
| | Fecundity–RWb-5 | 85 | 0.06 | 0.120 | 0.148 | 0.457 | 0.46 | 0.876 | 0.013 |
| | Mt Lewis | 40 | 0.32 | 0.640 | 0.148 | 0.457 | 2.46 | 0.490 | - |
| | Paluma | 45 | -0.21 | -0.420 | 0.148 | 0.457 | -1.61 | 0.869 | - |
| | RWb-5–Fecundity | 82 | -0.04 | -0.080 | 0.457 | 0.148 | -0.31 | 0.932 | 0.658 |
| | Fecundity–RWb-6 | 85 | -0.19 | -0.380 | 0.148 | 0.035 | -5.28 | 0.655 | 0.683 |
| | RWb-6–Fecundity | 82 | -0.13 | -0.260 | 0.035 | 0.148 | -3.61 | 0.869 | 0.592 |
| | Wing Size–RWb-1 | 81 | 0.08 | 0.160 | 0.476 | 0.462 | 0.34 | 0.869 | 0.759 |
| | RWb-1–Wing Size | 81 | -0.04 | -0.080 | 0.462 | 0.476 | -0.17 | 0.932 | 0.293 |
| | Wing Size–RWb-2 | 81 | 0.12 | 0.240 | 0.476 | 0.356 | 0.58 | 0.869 | 0.218 |
| | RWb-2–Wing Size | 81 | 0.07 | 0.140 | 0.356 | 0.476 | 0.34 | 0.869 | 0.327 |
| | Wing Size–RWb-3 | 81 | -0.07 | -0.140 | 0.476 | 0.680 | -0.25 | 0.869 | 0.579 |
| | RWb-3–Wing Size | 81 | 0.07 | 0.140 | 0.680 | 0.476 | 0.25 | 0.869 | 0.980 |
| | Wing Size–RWb-4 | 81 | -0.03 | -0.060 | 0.476 | 0.284 | -0.16 | 0.932 | 0.734 |
| | RWb-4–Wing Size | 81 | 0.03 | 0.060 | 0.284 | 0.476 | 0.16 | 0.932 | 0.156 |
| | Wing Size–RWb-5 | 81 | 0.07 | 0.140 | 0.476 | 0.457 | 0.30 | 0.869 | 0.297 |
| | RWb-5–Wing Size | 81 | -0.08 | -0.160 | 0.457 | 0.476 | -0.34 | 0.869 | 0.993 |
| | Wing Size–RWb-6 | 81 | -0.03 | -0.060 | 0.476 | 0.035 | -0.46 | 0.932 | 0.663 |
| | RWb-6–Wing Size | 81 | 0.02 | 0.040 | 0.035 | 0.476 | 0.31 | 0.960 | 0.643 |

| Species | Dam - daughter Population | Benign (23ºC) | | | | | | | |
|---|---|---|---|---|---|---|---|---|---|
| | | *N* | $\beta$ | $cov_{XY}$ | $cov_{XX}$ | $cov_{YY}$ | $r_G$ | *P*-value | *P*-value$_{pop}$ |
| *D. serrata* | Fecundity–Wing size | 63 | -0.08 | -0.160 | 0.052 | 1.000 | -0.70 | 0.932 | 0.292 |
| | Wing size–Fecundity | 70 | -0.16 | -0.320 | 1.000 | 0.052 | -1.40 | 0.869 | 0.078 |
| | Fecundity–RWs-1 | 63 | 0.11 | 0.220 | 0.052 | 1.000 | 0.96 | 0.869 | 0.736 |
| | RWs-1–Fecundity | 86 | 0.06 | 0.120 | 1.000 | 0.052 | 0.53 | 0.932 | 0.821 |
| | Fecundity–RWs-2 | 63 | -0.09 | -0.180 | 0.052 | 0.836 | -0.86 | 0.869 | 0.832 |
| | RWs-2–Fecundity | 68 | 0.08 | 0.160 | 0.836 | 0.052 | 0.77 | 0.869 | 0.001 |
| | Paluma | 34 | 0.42 | 0.840 | 0.836 | 0.052 | 4.03 | 0.234 | - |
| | Raglan Ck | 34 | -0.36 | -0.720 | 0.836 | 0.052 | -3.45 | 0.455 | - |
| | Fecundity–RWs-3 | 63 | 0.1 | 0.200 | 0.052 | 1.000 | 0.88 | 0.869 | 0.857 |
| | RWs-3–Fecundity | 68 | 0.29 | 0.580 | 1.000 | 0.052 | 2.54 | 0.234 | 0.905 |
| | Fecundity–RWs-4 | 63 | -0.12 | -0.240 | 0.052 | 0.514 | -1.47 | 0.869 | 0.934 |
| | RWs-4–Fecundity | 68 | -0.03 | -0.060 | 0.514 | 0.052 | -0.37 | 0.932 | 0.469 |
| | Fecundity–RWs-5 | 63 | 0.11 | 0.220 | 0.052 | 0.514 | 1.35 | 0.869 | 0.511 |
| | RWs-5–Fecundity | 68 | 0.12 | 0.240 | 0.052 | 0.510 | 1.47 | 0.869 | 0.106 |
| | Wing Size–RWs-1 | 63 | 0.01 | 0.020 | 1.000 | 1.000 | 0.02 | 0.981 | 0.630 |
| | RWs-1–Wing Size | 62 | 0.17 | 0.340 | 1.000 | 1.000 | 0.34 | 0.869 | 0.364 |
| | Wing Size–RWs-2 | 63 | 0.19 | 0.380 | 1.000 | 0.836 | 0.42 | 0.869 | 0.771 |
| | RWs-2–Wing Size | 62 | -0.04 | -0.080 | 0.836 | 1.000 | -0.09 | 0.932 | 0.631 |
| | Wing Size–RWs-3 | 63 | 0.08 | 0.160 | 1.000 | 1.000 | 0.16 | 0.869 | 0.421 |
| | RWs-3–Wing Size | 62 | 0.13 | 0.260 | 1.000 | 1.000 | 0.26 | 0.869 | 0.667 |
| | Wing Size–RWs-4 | 63 | 0.17 | 0.340 | 1.000 | 0.514 | 0.47 | 0.869 | 0.896 |
| | RWs-4–Wing Size | 62 | 0.01 | 0.020 | 0.514 | 1.000 | 0.03 | 0.981 | 0.296 |
| | Wing Size–RWs-5 | 63 | -0.15 | -0.300 | 1.000 | 0.510 | -0.42 | 0.869 | 0.076 |
| | RWs-5–Wing Size | 62 | 0.1 | 0.200 | 0.510 | 1.000 | 0.28 | 0.869 | 0.790 |

| Species | Dam - daughter Population | Stressful (28 ºC) | | | | | | | |
|---|---|---|---|---|---|---|---|---|---|
| | | *N* | *β* | $cov_{XY}$ | $cov_{XX}$ | $cov_{YY}$ | $r_G$ | *P*-value | *P*-value$_{pop}$ |
| *D. serrata* | Fecundity–Wing size | 63 | 0.02 | 0.040 | 0.040 | 0.226 | 0.42 | 0.981 | 0.715 |
| | Wing size–Fecundity | 64 | 0.06 | 0.120 | 0.226 | 0.040 | 1.26 | 0.944 | 0.769 |
| | Fecundity–RWs-1 | 63 | 0.05 | 0.100 | 0.040 | 0.268 | 0.97 | 0.932 | 0.609 |
| | RWs-1–Fecundity | 64 | -0.02 | -0.040 | 0.268 | 0.040 | -0.39 | 0.960 | 0.641 |
| | Fecundity–RWs-2 | 63 | 0.12 | 0.240 | 0.040 | 0.668 | 1.47 | 0.869 | 0.149 |
| | RWs-2–Fecundity | 64 | 0.08 | 0.160 | 0.668 | 0.040 | 0.98 | 0.869 | 0.217 |
| | Fecundity–RWs-3 | 63 | -0.01 | -0.020 | 0.040 | 0.576 | -0.13 | 0.981 | 0.504 |
| | RWs-3–Fecundity | 64 | -0.08 | -0.160 | 0.576 | 0.040 | -1.05 | 0.869 | 0.850 |
| | Fecundity–RWs-4 | 63 | 0.36 | 0.720 | 0.040 | 0.414 | 5.60 | 0.164 | 0.057 |
| | RWs-4–Fecundity | 64 | -0.22 | -0.440 | 0.414 | 0.040 | -3.42 | 0.651 | 0.331 |
| | Paluma | 26 | 0.05 | 0.100 | 0.414 | 0.040 | 0.78 | 0.932 | - |
| | Raglan Ck | 38 | -0.46 | -0.920 | 0.414 | 0.040 | -7.15 | 0.164 | - |
| | Fecundity–RWs-5 | 63 | 0.01 | 0.020 | 0.040 | 0.430 | 0.15 | 0.981 | 0.183 |
| | RWs-5–Fecundity | 64 | 0.00 | 0.000 | 0.430 | 0.040 | 0.00 | 0.983 | 0.293 |
| | Wing Size–RWs-1 | 62 | -0.31 | -0.620 | 0.226 | 0.268 | -2.52 | 0.234 | 0.419 |
| | RWs-1–Wing Size | 62 | -0.09 | -0.180 | 0.268 | 0.226 | -0.73 | 0.869 | 0.748 |
| | Wing Size–RWs-2 | 62 | 0.05 | 0.100 | 0.226 | 0.668 | 0.26 | 0.932 | 0.518 |
| | RWs-2–Wing Size | 62 | -0.06 | -0.120 | 0.668 | 0.226 | -0.31 | 0.932 | 0.171 |
| | Wing Size–RWs-3 | 62 | -0.13 | -0.260 | 0.226 | 0.576 | -0.72 | 0.869 | 0.908 |
| | RWs-3–Wing Size | 62 | 0.11 | 0.220 | 0.576 | 0.226 | 0.61 | 0.869 | 0.823 |
| | Wing Size–RWs-4 | 62 | 0.07 | 0.140 | 0.226 | 0.414 | 0.46 | 0.876 | 0.035 |
| | Paluma | 26 | 0.38 | 0.760 | 0.226 | 0.414 | 2.48 | 0.512 | - |
| | Raglan Ck | 36 | -0.16 | -0.320 | 0.226 | 0.414 | -1.05 | 0.869 | - |
| | RWs-4–Wing Size | 62 | -0.10 | -0.200 | 0.414 | 0.226 | -0.65 | 0.869 | 0.366 |
| | Wing Size–RWs-5 | 62 | 0.18 | 0.360 | 0.226 | 0.430 | 1.15 | 0.869 | 0.227 |
| | RWs-5–Wing Size | 62 | 0.10 | 0.200 | 0.430 | 0.226 | 0.64 | 0.869 | 0.917 |

**Supplementary Table 6**: **Genetic correlations between fitness, wing size, and wing shape in two different thermal environments**.

$r_G$ denotes the genetic correlation between trait one ($trait_1$) and trait two ($trait_2$) and *SE* is the standard error for the genetic correlation. Wing shape variables are shown as the relative warp (RW) scores that contribute to > 5% variability. Bold values indicate a statistically significant correlation based on standard errors. *P*-values were adjusted for False Discovery Rate and significance level is shown with asterisks (**P* < 0.05, ***P* < 0.01, ****P* < 0.001, *****P* < 0.0001).

| | | | Benign (23ºC) | | | Stressful (28 ºC) | | |
|---|---|---|---|---|---|---|---|---|
| Species | $trait_1$ | $trait_2$ | $r_G$ | *SE* | *P*-value | $r_G$ | *SE* | *P*-value |
| *D. birchii* | Fecundity | Wing size | 0.19 | 0.32 | 1.000 | - | - | - |
| | Fecundity | RWb-1 | -0.15 | 0.11 | 1.000 | - | - | - |
| | Fecundity | RWb-2 | 0.17 | 0.28 | 1.000 | - | - | - |
| | Fecundity | RWb-3 | -0.32 | 0.19 | 1.000 | - | - | - |
| | Fecundity | RWb-4 | -0.24 | 0.39 | 1.000 | - | - | - |
| | Fecundity | RWb-5 | 0.08 | 0.29 | 1.000 | - | - | - |
| | Fecundity | RWb-6 | 0.00 | 0.00 | 0.000 | - | - | - |
| | Wing size | RWb-1 | 0.09 | 0.07 | 0.500 | - | - | - |
| | Wing size | RWb-2 | **0.46** | 0.14 | 0.006** | - | - | - |
| | Wing size | RWb-3 | 0.00 | 0.13 | 1.000 | - | - | - |
| | Wing size | RWb-4 | 0.00 | 0.25 | 1.000 | - | - | - |
| | Wing size | RWb-5 | -0.02 | 0.17 | 1.000 | - | - | - |
| | Wing size | RWb-6 | -0.08 | 0.21 | 1.000 | - | - | - |
| *D. serrata* | Fecundity | Wing size | **-1.00** | 0.04 | 6.96E-120**** | **0.84** | 0.29 | 0.019* |
| | Fecundity | RWs-1 | **0.75** | 0.16 | 2.60E-05**** | 0.29 | 0.76 | 1.000 |
| | Fecundity | RWs-2 | -0.05 | 0.45 | 1.000 | 1.00 | 0.27 | 0.166 |
| | Fecundity | RWs-3 | 1.00 | 0.73 | 1.000 | -0.59 | 0.43 | 1.000 |
| | Fecundity | RWs-4 | **-0.92** | 0.10 | 4.37E-12**** | **1.00** | 0.13 | 4.44E-08**** |
| | Fecundity | RWs-5 | 1.00 | 0.49 | 1.000 | 0.08 | 0.71 | 1.000 |
| | Wing size | RWs-1 | 0.18 | 0.08 | 0.094 | -1.00 | 0.54 | 0.203 |
| | Wing size | RWs-2 | 0.16 | 0.11 | 0.394 | -0.03 | 0.22 | 1.000 |
| | Wing size | RWs-3 | 0.21 | 0.09 | 0.085 | -0.06 | 0.27 | 1.000 |
| | Wing size | RWs-4 | 0.25 | 0.14 | 0.215 | -0.10 | 0.28 | 1.000 |
| | Wing size | RWs-5 | -0.07 | 0.12 | 1.000 | **0.90** | 0.06 | 8.36E-45**** |

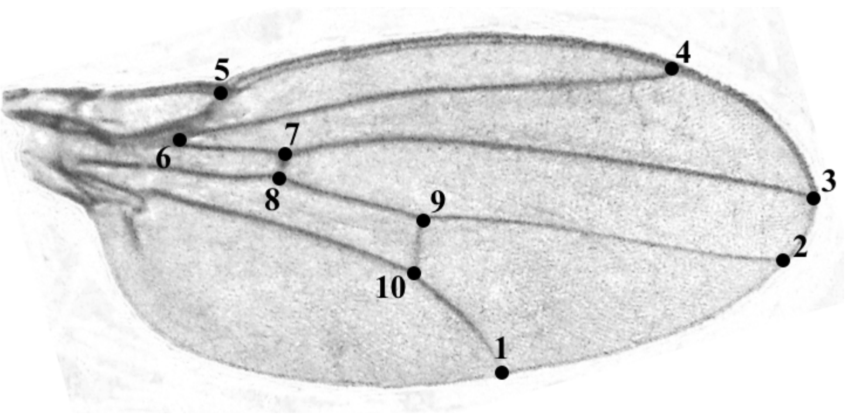

**Supplementary Figure 1: Landmarks for wing size and wing shape measurements.**

An example of a wing image, showing the 10 landmark positions in sequential order that were used to compute centroid size for wing size and relative warp scores for wing shape.

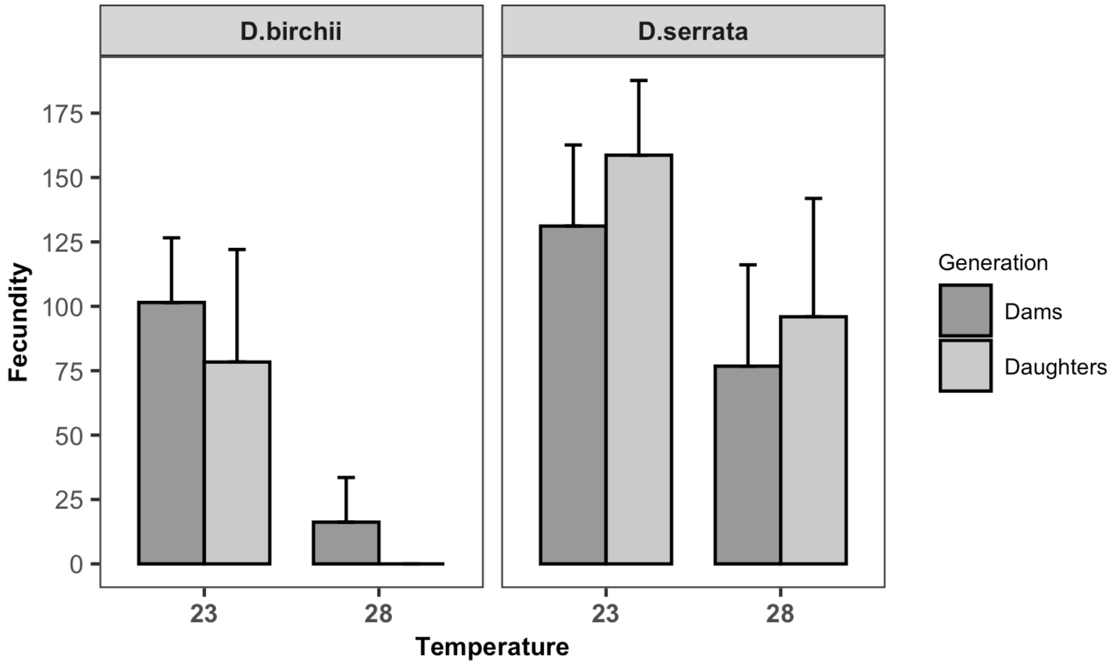

**Supplementary Figure 2: Fecundity of dams and their daughters exposed to two different thermal environments for the entirety of their life**.

Fecundity is based on total egg count of 72 hrs. Error bars show the standard deviations and means and sample sizes are shown in Supplementary Table 1.

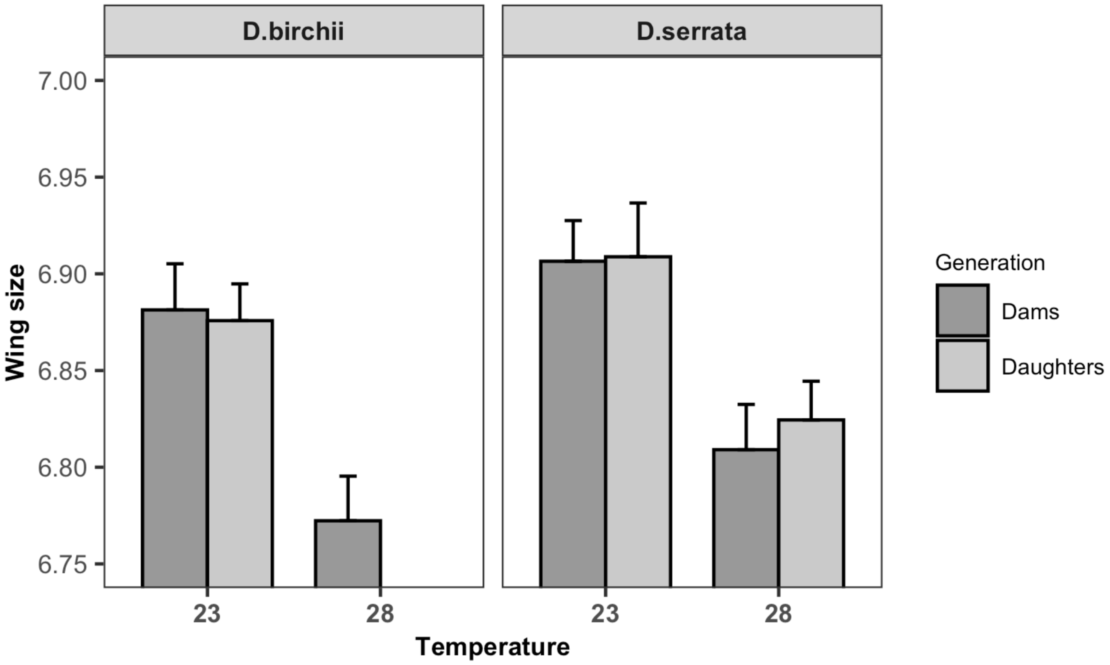

**Supplementary Figure 3: Wing size of dams and their daughters exposed to two different thermal environments for the entirety of their life.**

Wing size is shown as log centroid size that is measured in arbitrary units. Error bars show the standard deviations and means and sample sizes are shown in Supplementary Table 2.

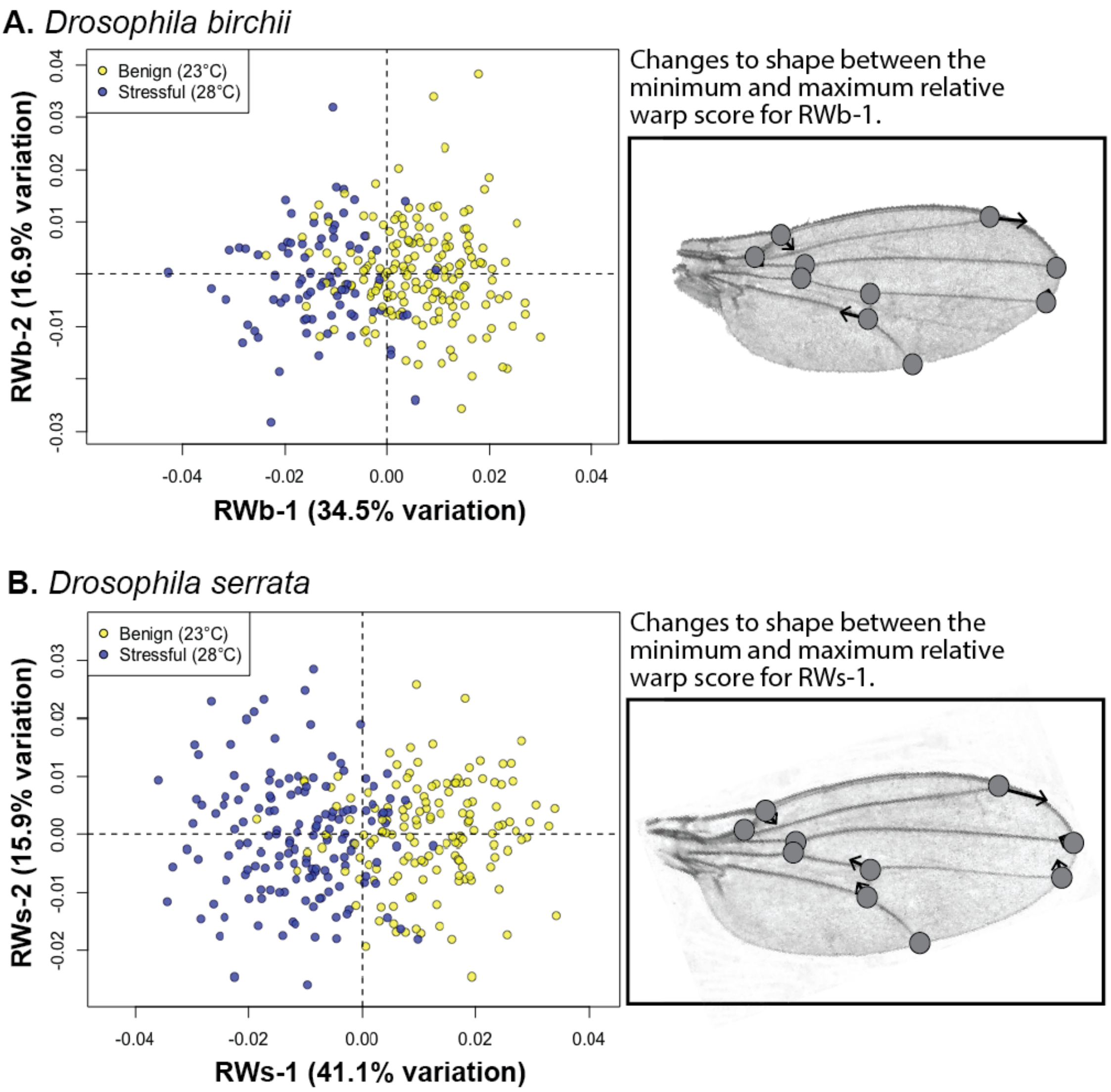

**Supplementary Figure 4: PCA plots showing the RW 1 and RW 2 axes for wing shape in (A) *Drosophila birchii* and (B) *D. serrata* grouped by temperature treatment and for all individuals measured (i.e., both dams and daughters).**

Enlarged images of wings show the directionality and position (indicated by the black arrows) of change to each landmark between the minimum (shown) and maximum (end of the arrow) relative warp score for RWb/s-1. Means and samples sizes are shown in Supplementary Table 3.
